# Anxiolytic treatment impairs helping behavior in rats

**DOI:** 10.1101/044180

**Authors:** I. Ben-Ami Bartal, H. Z. Shan, N. M. R. Molasky, T. M. Murray, J. Z. Williams, J. Decety, Peggy Mason

## Abstract

Despite decades of research with humans, the biological mechanisms that motivate an individual to help others remain poorly understood. In order to investigate the roots of pro-sociality in mammals, we established the helping behavior test, a paradigm in which rats are faced with a conspecific trapped in a restrainer that can only be opened from the outside. Over the course of repeated test sessions, rats exposed to a trapped cagemate learn to open the door to the restrainer, thereby helping the trapped rat to escape (Ben-Ami Bartal et al., 2011). The discovery of this natural behavior provides a unique opportunity to probe the motivation of rodent helping behavior, leading to a deeper understanding of biological influences on human pro-sociality.

To determine if an affective response motivates door-opening, rats received midazolam, a benzodiazepine anxiolytic, and tested in the helping behavior test. Midazolam-treated rats showed less helping behavior than saline-treated rats or rats receiving no injection. Yet, midazolam-treated rats opened a restrainer containing chocolate, highlighting the socially specific effects of the anxiolytic. To determine if midazolam interferes with helping through a sympatholytic effect, the peripherally restricted beta-adrenergic receptor antagonist nadolol was administered; nadolol did not interfere with helping.

The corticosterone response of rats exposed to a trapped cagemate was measured and compared to the rats’ subsequent helping behavior. Rats with the greatest corticosterone responses showed the least helping behavior and those with the smallest responses showed the most consistent helping at the shortest latency. These results are discussed in terms of their implications for the interaction between stress and pro-social behavior.

Finally, we observed that door-opening appeared to be reinforcing. A novel analytical tool was designed to interrogate the pattern of door-opening for signs that a rat’s behavior on one session influenced his behavior on the next session. Results suggest that helping a trapped rat has a greater motivational value than does chocolate.

In sum, this series of experiments clearly demonstrates the fundamental role of affect in motivating pro-social behavior in rodents and the need for a helper to resonate with the affect of a victim.

## Introduction

Helping refers to actions that intentionally benefit others (Cronin, 2012). In humans, helping is often motivated by an empathic response to the distress and pain of others. Yet it is very hard to predict when helping will occur, and why some situations fail to elicit an empathic response. Moreover, several psychopathologies are marked by a lack of empathy and consequent detriments in pro-social behavior. To better understand help, it is imperative to parse out the biological mechanisms that give rise to an emotional response to the distress of others, a task that has seen only partial success with human studies. An animal model of pro-social behavior is crucial for investigating the biological cascade of events that occur in the moments between the observation of distress and the decision to act for the benefit of others.

Recently we established the rat helping paradigm in which a free rat is placed in an arena with a centrally located plastic tube (termed “restrainer”) containing a trapped rat (Ben-Ami Bartal et al., 2011). Over the course of repeated testing sessions, and with no external reward provided, free rats learn to open the restrainer and thereby release the trapped rat. In order to be considered a successful helper, or opener, the free rat needs to demonstrate consistent helping by releasing the trapped rat over several trials once learning to do so. Door-opening is effortful, demanding of exploratory behavior, and typically takes rats a few days to learn. While the trapped rat benefits from the free rat’s action, it remains debatable whether the free rat intends door-opening to serve the trapped rat’s benefit. The experiments here are designed to test if the free rat uses an affective motivation to open the restrainer door and thereby release the trapped rat.

There are several possible motivations for door-opening. A rat may open the restrainer door in response to a contextual clue unrelated to the trapped rat’s distress. For example, the rat may enjoy the motor mastery of opening the restrainer door although this is unlikely since rats do not open the doors to empty restrainers or object-containing restrainers (Ben-Ami Bartal et al., 2011). Another possibility is that rats open the door in order to play with the trapped rat or in order to terminate an aversive sensory cue, such as an alarm call or pheromone, emitted by the trapped rat. On the other hand, the free rat may catch the trapped rat’s distress and that socially acquired emotion may fuel the free rat’s door-opening act. Ample scientific evidence shows that “the affective feelings of one [rodent] are conveyed to another and then generate the same feelings in that individual” (reviewed in Panksepp & Lahvis 2011). In humans, such vicarious experience of distress often leads to helping. It has been unclear whether the same motivational mechanism drives helping in rodents. Therefore the present experiments are designed to test if rats must mount an affective response to the distress of a trapped rat in order to open the restrainer door and release the trapped rat. This would constitute a rodent form of empathy.

To test whether an affective state is required for the free rat to release a trapped cagemate, we treated free rats with midazolam (MDZ), a benzodiazepine anxiolytic. Midazolam, commonly used to treat anxiety in people, functions as a positive allosteric modulator of the GABA-A receptor. It thereby facilitates inhibition widely through its actions on the brain’s ubiquitous inhibitory ionotropic receptor. MDZ-treated rats were tested in the rat helping paradigm. Consistent with the hypothesis that the free rat is motivated by a vicarious experience of the trapped rat’s affect, MDZ-treated rats did not open the restrainer door for a trapped rat whereas control rats (uninjected, saline-treated) did.

As with all pharmacological agents, MDZ can have multiple, and in this case psychotropic, effects. In addition to serving as powerful anxiolytics, benzodiazepines are sedating; they reduce exploratory motor behavior and at high enough doses can serve as hypnotics (sleep-inducing drugs). Benzodiazepines can also reduce oxytocin transmission (Welt et al 2006, Yagi and Onaka 1996), which may be expected to impair social behavior (Anacker and Beery 2013, Febo and Ferris 2014). Other studies show that benzodiazepines promote approach and reduce aggression in rats (Weerts et al 1993, Christmas and Maxwell 1970). Finally, it is possible that benzodiazepines modify learning processes in some way. To control for any known, suspected, or unanticipated detrimental effect that MDZ may exert on a rat’s ability to learn to open a restrainer door, we tested the effect of MDZ treatment on door-opening to access chocolate, a non-social stimulus that elicits approach behavior and does not require social processing. We found that MDZ-treated rats readily opened the restrainer door to access chocolate, demonstrating that door-opening is not impaired by MDZ.

Finding that MDZ blocked door-opening for a trapped rat but not for chocolate, we wanted to further define the site of MDZ’s anti-empathic actions. Distress or anxiety can activate the hypothalamo-pituitary-adrenal (HPA) axis, which in turn results in increased sympathetic outflow. MDZ acts in the brain to reduce anxiety, which in turn reduces the downstream consequences of HPA and sympathetic activation. To distinguish between the direct and indirect actions of MDZ, the effects of nadolol, a peripherally acting beta-adrenergic receptor blocker that antagonizes sympathetic arousal in the rat, on door-opening were tested. There were no differences between the performance of nadolol-treated and control rats, evidence that the downstream (and indirect) sympathetic-dampening effects of MDZ are not responsible for MDZ’s anti-helping effects. Thus, MDZ appears to act either within the brain or on the HPA axis upstream of sympathetic activation.

To determine if antagonizing HPA reactions could account for MDZ’s detrimental effects on helping effects, we compared free rats’ HPA reaction to the trapped rat’s distress, as measured by corticosterone release, with the free rats’ subsequent door-opening behavior. In other words, free rats were exposed to a trapped rat (without the ability to release him) and their corticosterone response measured. On subsequent days, rats were then tested in the rat helping paradigm. The results show that there is a significant correlation between HPA reactivity to vicarious distress and door-opening latency, suggesting that HPA reactivity may antagonize helping (longer latencies). This possibility, along with several alternatives, is discussed in the Discussion.

We observed that after opening the restrainer, rats were highly likely to do so again in the subsequent test session. This pattern suggests that door-opening is reinforcing. We therefore designed a new analytical tool to interrogate the pattern of door-opening for signs that a rat’s behavior on one session influenced his behavior on the next session. If the probability of opening on one session depends on the outcome of the previous session, this would imply that in statistical terms, the door-opening pattern of a reinforced rat cannot be modeled by a rate-varying stochastic process. Importantly, increasing probability of opening across sessions alone is not evidence for reinforcement. Such an increase could be driven by more familiarity with contextual cues (restrainer, door, arena, timing of experiment, experimenters), higher motor proficiency, or decreased anxiety. These factors are not reinforcing in that they occur regardless of previous sessions’ outcomes and are therefore outcome-independent. Yet they still lead to increasing opening probabilities across sessions. To determine if outcome-independent effects can account for the observed opening patterns, we modeled door-openings with a rate-varying stochastic process and compared the simulated patterns to observed patterns as a stringent test of the outcome-dependence of door-opening.

## Material and methods

All procedures conformed to the established ethical standards and were reviewed and approved by the University of Chicago Institutional Animal Care and Use Committee.

*Subjects*. Two month-old Sprague-Dawley (SD) rats (Charles River, Portage, MI) were used for all studies. All rats were male and were housed in same-sex pairs. Rats had *ad libitum* access to chow and water in a 12:12 light-dark cycle, and were allowed two weeks to acclimate to the housing environment and their cagemate.

Three experiments were performed. Five groups of rats (n=16 rat pairs/group) were studied in the “basic paradigm” condition involving a trapped rat. Three groups of rats (n=8 rats/group) were studied in a modified condition with a chocolate-containing restrainer. Finally, 40 rats were studied for their corticosterone response to either being trapped (n=20) or to viewing their cagemate trapped (n=20) within the restrainer. After measuring the corticosterone response, these rats were tested in the rat helping paradigm with a trapped cagemate. Data from both trapped (n=20) and free (n=20) rats in this experiment are presented.

In sum, a total of 114 test rats were studied in the trapped and chocolate conditions. An additional 40 rats (20 pairs) were studied for their corticosterone response to either being trapped (n=20) or to viewing their cagemate trapped (n=20). No dropouts occurred.

*Habituation*. Two weeks after arriving at the animal facility, animals were habituated to the testing rooms, experimenters (who were kept constant for each cohort of rats), and testing arenas. Testing arenas were constructed of Plexiglas (50 × 50 cm, 32-60 cm high) and were kept constant for each pair or rats. On day 1 of habituation, rats were transported to the testing room and left undisturbed in their home cages. Thereafter, rats were transported to the room and left undisturbed for 15 min prior to habituation procedures. On day 2, rats were briefly handled. Starting with the second day of habituation, rats were weighed 3 times weekly for the duration of the experiment; no animal lost weight during the experiment. On days 3-6, rats were handled for 5 minutes by each experimenter and then placed together (in housing pairs) in the testing arenas for 30 minutes. After each habituation session, rats were returned to their home cages and to the housing room. Within each cage, rats were randomly chosen to be either the free or trapped rat. Rats did not switch roles.

In order to habituate free rats to i.p. injections and minimize stress related to the injection itself, free rats in injection groups (4 groups of 16 each in the trapped condition, 3 groups of 8 each in the chocolate condition) received i.p. saline injections once daily for at least 5 days preceding testing. After receiving these saline injections, rats were placed in the testing arenas for 30 minutes as above. Rats in the uninjected (n=16) and corticosterone (n=10 pairs) groups received no injections during habituation.

*Open field testing*. On the day following completion of habituation, rats were placed individually in an arena for 30 min and their activity recorded. The arenas were the same as were used during habituation but that open field testing represented the first time each rat had been in the arena alone. Open field testing has been done routinely as a minimally invasive metric of individual rat behavior. As it turns out, data from open field testing are not included in this report. Nonetheless, the animals experienced this testing and we therefore include it to provide a complete account of the rats’ treatment.

*Protocol for trapped and chocolate conditions*. On each testing day, rats were transported to the testing room and left undisturbed in their home cage for 15 minutes. Then rats were colored with markers to permit tracking the rats’ individual movements. The free rat was colored red and the trapped rat colored blue. Rats were then weighed after coloring.

After coloring and weighing, rats in the uninjected group were placed into the arenas for testing. Rats in injection groups received MDZ (2 mg/kg for the high dose conditions; 1.25 mg/kg for the low dose conditions, i.p.), nadolol (10 mg/kg), or saline (0.5 cc, i.p.). As explained in the Introduction, MDZ is a benzodiazepine that acts on the brain to produce anxiolytic and sedative effects. Nadolol is a beta-adrenergic antagonist that does not cross the blood brain barrier; it blocks sympathetic effectors but not corticosterone release and does not have central anxiolytic effects. Saline is a vehicle control.

After the free rats received an injection, they were returned to their home cage. After a waiting period (15 min for MDZ and saline; 30 min for nadolol), rats were placed in the arena and the helping behavior test began.

*Trapped rat paradigm*. The trapped rat was placed inside a restrainer and the restrainer was positioned in the arena center. Restrainers were Plexiglas tubes (25 × 8.75 × 7.5 cm; Harvard Apparatus, Holliston, MA) that had several slits, allowing for olfactory, auditory, and tactile communication between rats. The free rat (the trapped rat’s cagemate) was then placed in the arena and allowed to roam freely. The door to the restrainer could only be opened from the outside and therefore only by the free rat. If the free rat did not open the restrainer door within 40 min, the investigator opened the restrainer door “halfway,” to a 45° angle, greatly facilitating door-opening by either rat. Only door-openings that occurred prior to the halfway opening were counted as such.

Rat dyads always remained in the arena for a full hour. Hour-long testing sessions were repeated for 12 days and performed only once per day. All sessions were run during the rats' light cycle between 0800 and 1730. After each session, rats were returned to their home cages and the arena and restrainer were washed with 1% acetic acid followed by surface cleaner.

*Blockers*. Some trapped rats (n=30, 38%) succeeded in opening the door from inside the restrainer during one of the testing sessions. When this happened, the trapped rat was placed immediately back in the restrainer, and a Plexiglas blocker was inserted, preventing his access to the door. If the free rat subsequently opened the door, the blocker was removed, allowing the trapped rat to exit the restrainer. The blocker was then used for that trapped rat on all following test days. If the free rat failed to open the door by 40 min, the blocker was removed when the door was opened halfway.

*Chocolate condition*. Rats in the three chocolate conditions (high and low MDZ, saline), were introduced to chocolate chips prior to the experimental sessions. After this exposure, they ate an average of 4.6 ± 0.4 chips at a time. On testing days, the restrainer was filled with 5 chocolate chips (Nestlé® Toll House, milk chocolate) and positioned in the arena center; chocolate was not available to rats outside of the testing sessions. The free rat was placed in the arena with the restrainer but without his cagemate; all other details of the experimental protocol were as described above. When rats opened the restrainer door, they always ate all 5 chips.

*Door-opening analysis*. Latency to door-opening was calculated as the minute when the restrainer door was opened minus the start time. For rats that never opened, a cutoff time of 40 min (the time of halfway opening) was assigned.

*Corticosterone measurements*. Blood samples were collected via tail-nick from a cohort of 40 male rats housed in 20 pairs. Rats were habituated to the arenas for 10 days. Then rats were placed in the arenas with one rat trapped and one rat free (roles chosen at random). The restrainer was taped shut on this testing day, in order to ensure that all free rats were exposed to a trapped rat for a full 40 minutes. Blood was then collected to determine each rat’s corticosterone response to either being trapped (n=20) or to witnessing a cagemate being trapped (n=20).

Blood collection occurred at three time points. The first sample (baseline) was collected an hour prior to placement in the arena. The second sample (test) was collected immediately after removal from the arena, and the third sample (post) was collected an hour after removal from the arena. Pilot experiments revealed that unstressed, male Sprague-Dawley rats show steady levels of CORT between 0830 and 1230. Therefore all samples were collected during this time period.

Blood (200-500 μL) was collected via tail-nick, by experimenters who had handled the rats previously. Sampling was completed in less than three minutes (average 2:18). Samples were immediately centrifuged for 10 min at 5000 rpm at 4°C. Plasma was extracted and frozen at −20°C for further analysis via enzyme-linked immunoabsorbent assay (ELISA, IBL). The assay had a sensitivity of < 27.0 pg/ml. Two outliers were removed from analysis.

*Behavioral testing following corticosterone measurement*. After the day of blood collection, the 20 pairs of male rats were tested in the basic trapped rat paradigm described above for 12 days.

*Statistical analysis of opening latency*. For the trapped and chocolate experiments, opening latencies of each subject (16 per trapped group, 8 per chocolate group) on each day (12 per subject) from each experimental group (5 trapped groups, 3 chocolate groups) were analyzed using a general linear model with the statistical software *R (R Foundation for Statistical Computing, Vienna, Austria, used under the General Public License)* and the R package "regress" *(David Clifford and Peter McCullagh, used under the General Public License)*. The R code for the analysis as well as the original latency data (Supplement B) are included below.

Although a repeated-measure (time) two-level (drug, rat) design is traditionally analyzed using a two-way repeated-measure ANOVA, we used a General Linear Model (GLM) for reasons that are fully explained in Appendix A. In brief, only a GLM can account for differences in the correlation between two data points, separated by different time intervals, from the same subject. For example, in an ANOVA, the latencies from day 1 and day 2 are expected to correlate to each other to the same extent as latencies from day 1 and day 8. However this is not the case in an appropriately crafted GLM. In the present GLM, *Vrat* was constructed as a correlation matrix that informs the model which data points are from the same subjects (repeated measure), and how the within-subject correlation decays with increasing intervals time (the correlation between latencies on days n and n+1 is greater than the correlation between latencies on days n and n+5 for example).

A second advantage of using GLM over an ANOVA is that we can directly test hypotheses, rather than relying on post-hoc tests. For each experiment, we built two models, one with the factors listed above (alternative model) and another with all the factors listed except interaction between treatment and day (null model). We fitted the data linearly onto these two models, and compare their goodness-of-fit to decide whether the interaction significantly improves the explanatory power of the model.

*Statistical analysis of opening results (binary) to test for reinforced behaviors*. While there were significantly different numbers of openings from rats that received different drug treatments, it is unclear whether this difference stems from different levels of reinforcement (after having opened, some rats are more inclined to open again on the next session) or differences in when rats try opening for the first time, perhaps related to anxiety, familiarity, or perceptual learning. Clearly, the former is a better metric of “willingness” to open. In sum, reinforcement would serve to make rats more likely to open sequentially (opening on consecutive days) than would random exploration or other non-reinforced behaviors.

To test whether rats were reinforced to open, or they merely opened by chance as they explored in the arena, we built a stringent null model that shows what the opening pattern would be if rats were not reinforced, but still opened at the same pace. Taking a leaf out of the playbook of estimating neurons’ spiking rates, we first calculated the opening probability of each rat on each day (see Supplement C). We then simulated rats’ opening by using a binary process to decide whether each rat opens on any given day. Our simulation successfully reproduces the total number of openings in different treatment groups, as well as their overall structure across time and rats (i.e. some rats open and some don’t; rats that open, open more on later days than earlier days). We then ran the simulation repeatedly, calculating the probability of Sequential Opening (%SO) of each of the null populations. Finally we determined if observed rats opened sequentially significantly more than did null rats.

## Results

### Overall design

Three experiments are reported. In the first experiment, five groups of rats (n=16 rats/group) were tested with a trapped cagemate. Rats were either not injected or injected with saline, a low or high dose of the benzodiazepine MDZ, or nadolol, a beta-adrenergic receptor antagonist that does not cross the blood-brain barrier (see Introduction for rationales). Comparisons of opening behavior were made between the five groups.

In the second experiment, three groups of rats (n=8 rats/group) were studied in a modified setup with a chocolate-containing restrainer. Rats in these conditions received injections of either saline, a low dose of MDZ, or a high dose of MDZ. Comparisons of opening behavior were made between the groups.

In a final experiment, 20 pairs of rats were studied for their corticosterone response to either being trapped (n=20) or to viewing their cagemate trapped (n=20). During this exposure to direct (being trapped) or vicarious (viewing the cagemate being trapped) stress, the restrainer restrainer could not be opened. After measuring the corticosterone response to direct or vicarious stress, rats were tested in the basic paradigm over the subsequent 12 days. A within-subjects comparison between the corticosterone response measured and the mean door-opening latency during subsequent testing was performed.

### Blocking distress in free rats tested with a trapped cagemate

Rats that received no injection were compared to rats that received either vehicle (saline) or one of two doses (low 1.25 mg/kg; high 2.0 mg/kg) of the benzodiazepine anxiolytic, MDZ. To distinguish between the direct anxiolytic effects and secondary and peripheral sympathetic- dampening effects of MDZ, a group of rats received nadolol, a beta-adrenergic antagonist that does not cross the blood-brain barrier.

Overall, the opening latency decreased across days, reflecting learning (Fig. 1A). This decay in opening latency across days differed between treatment groups (general linear model as described in the Methods: χ^2^(4)=12.0; p=0.02; Fig. 1B; Table 1). Untreated rats as well as rats treated with saline or nadolol showed decreasing opening latencies across the days of testing (linear model analysis; uninjected: N(0,1)=−4.36, p<0.001; saline: N(0,1)=−3.56, p<0.001; nadolol: N(0,1)=−3.86, p<0.001). In contrast, there was no decay in latency across days for rats treated with either dose of MDZ (linear model analysis; low: N(0,1)=−1.67, p=0.09; high: N(0,1)=−0.19, p=0.85). Thus, rats treated with MDZ did not show evidence of learning across the test sessions. Interestingly, the average opening latency of rats treated with the high dose of MDZ started high and remained high throughout testing whereas the latency of rats treated with the low dose of MDZ tended to be low on the initial days of testing and relatively high on the final days of testing, giving rise to a shallow and non-significant downward trend in latency (p=0.09).

**Figure 1.**
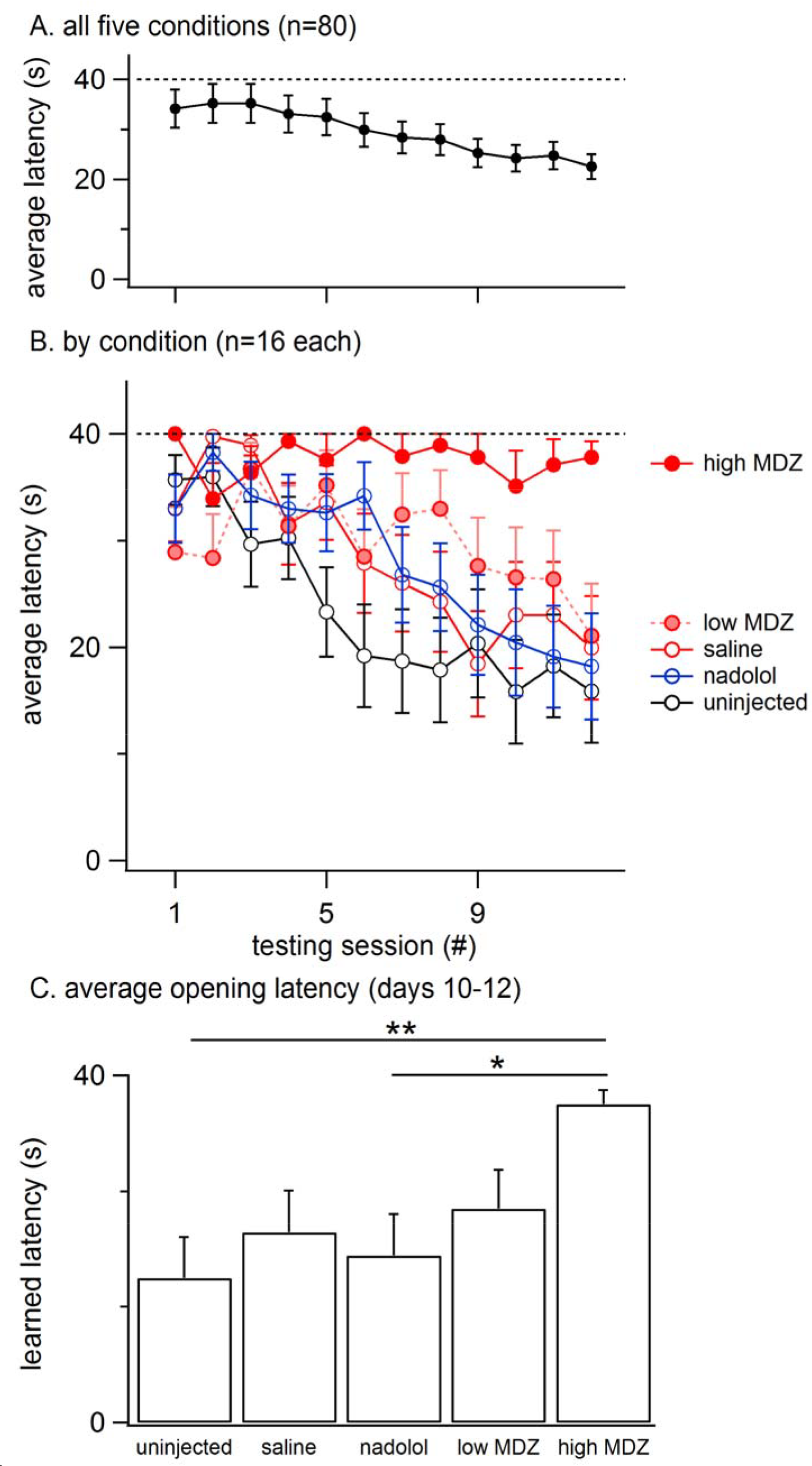
A: The mean (± SEM) latency to door-opening for all rats (n=80) decreased across the 12 days of testing, suggestive of learning. B: The decay in door-opening latency across testing sessions differed between the groups of rats tested (n=16 per group). C: The average opening latency during the final 3 days of testing, when latencies had plateaued, was significantly greater for rats treated with a high dose of MDZ than for rats that received no injection (**, p=0.01) or an injection of nadolol (*, p=0.04).

**Table 1.** Opening latencies declined across days. The decay of opening latency in each condition was compared to the decay observed for every other condition. P-values for these pair-wise comparisons are displayed (see Supplement for code).

|  | Uninjected | Saline | Nadolol | Low MDZ |
| --- | --- | --- | --- | --- |
| Uninjected |  |  |  |  |
| Saline | 0.58 |  |  |  |
| Nadolol | 0.73 | 0.83 |  |  |
| Low MDZ | 0.06 | 0.18 | 0.12 |  |
| High MDZ | <0.01 * | 0.02 * | <0.01 * | 0.29 |

The most pronounced drops in latency occurred during the testing sessions on the middle 5-6 days (Fig. 1A). Therefore, to examine rats’ stabilized performance, rather than the learning rate, the average latency recorded during days 10-12 was calculated; this was termed the *learned latency*. The learned latency was different between groups (Fig. 1C; one-way ANOVA; F(4, 75)=3.315, p=0.02). The learned latency of rats treated with a high dose of MDZ was significantly greater than that of uninjected rats (Tukey post hoc, p=0.01) or rats treated with nadolol (Tukey post hoc, p=0.04).

The group averages shown in Figure 1B–Figure 1C and Table 2 fail to reveal an important within-group variation that resulted from two subpopulations of animals. In each condition, at least six rats, and as many as ten, never consistently opened the restrainer with some of these rats never opening the restrainer at all. Figure 2A reveals the two different subpopulations in each of the five conditions studied. Box plots show the downward trend in the median value (for the non-MDZ-treated groups) as well as the shift of the latency distribution between days 1, 6 and 12 of testing (blue vertical histograms on right). In the low MDZ condition, a modest shift in opening latency distribution was observed in a subset of rats. In the high MDZ condition, no shift in opening latency distribution was observed.

**Table 2.** Mean (± SEM) door-opening latency and number of door-openings for all conditions.

|  | Mean latency | Mean number of openings<br>/12 days |
| --- | --- | --- |
| <u>Trapped conditions</u> |  |  |
| Uninjected | $23.4 \pm 1.9$ | $5.7 \pm 1.2$ |
| Saline | $28.4 \pm 1.8$ | $4.0 \pm 0.9$ |
| Nadolol | $28.1 \pm 1.7$ | $4.3 \pm 0.8$ |
| Low MDZ | $29.7 \pm 1.7$ | $3.9 \pm 0.9$ |
| High MDZ | $37.7 \pm 0.8$ | $1.0 \pm 0.5$ |
| <u>Chocolate conditions</u> |  |  |
| Saline | $34.9 \pm 1.8$ | $2.0 \pm 1.3$ |
| Low MDZ | $19.6 \pm 2.7$ | $7.0 \pm 1.5$ |
| High MDZ | $25.3 \pm 2.7$ | $5.0 \pm 1.5$ |

**Figure 2.**
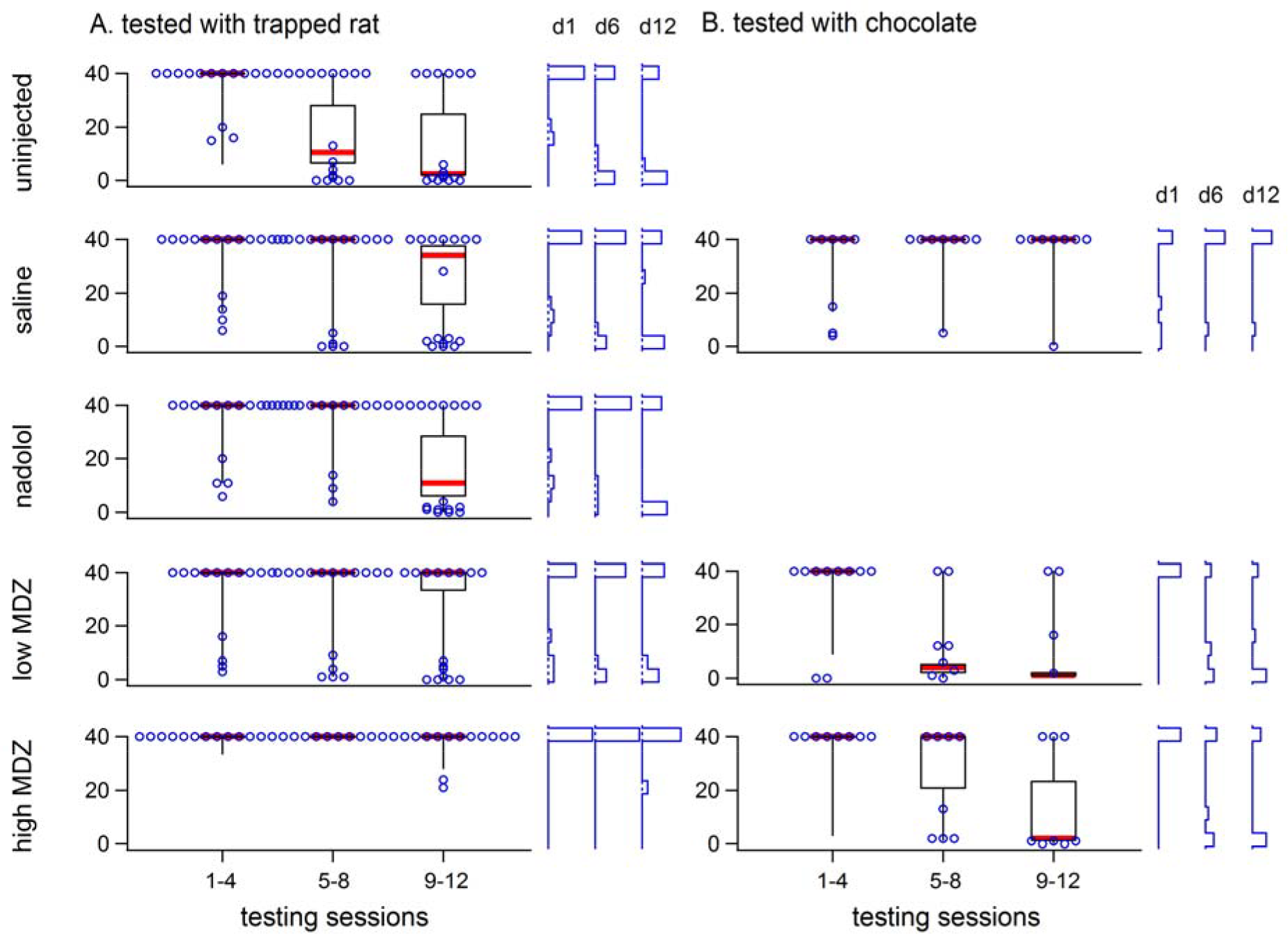
A: The variability of opening latency within groups is illustrated using box plots (40, 50, 60 percentile lines with the median marked in red, 10 and 90 percentile whiskers) showing latencies across the 12 days of testing. All individual latencies are ilustrated for days 1, 6, and 12 (hollow blue circles). Frequency histograms of latencies on those days are shown at the right for each group. In all groups except the high MDZ rats tested with a trapped rat and saline rats tested with chocolate, there was a shift from long to short latencies.

## Testing for reinforcement

As detailed in the methods and appendices, we constructed a null model that estimates the proportion of openings (day 1-11) that would be followed by another opening (%SO) in the absence of reinforcement from one session to the next session. We then compared observed %SO values to null %SO values. The distribution of null %SO values for all randomly generated matrices are illustrated in Figure 3 along with the two-tailed probability that the observed %SO (red dotted line) came from the null distribution of %SO values.

**Figure 3.**
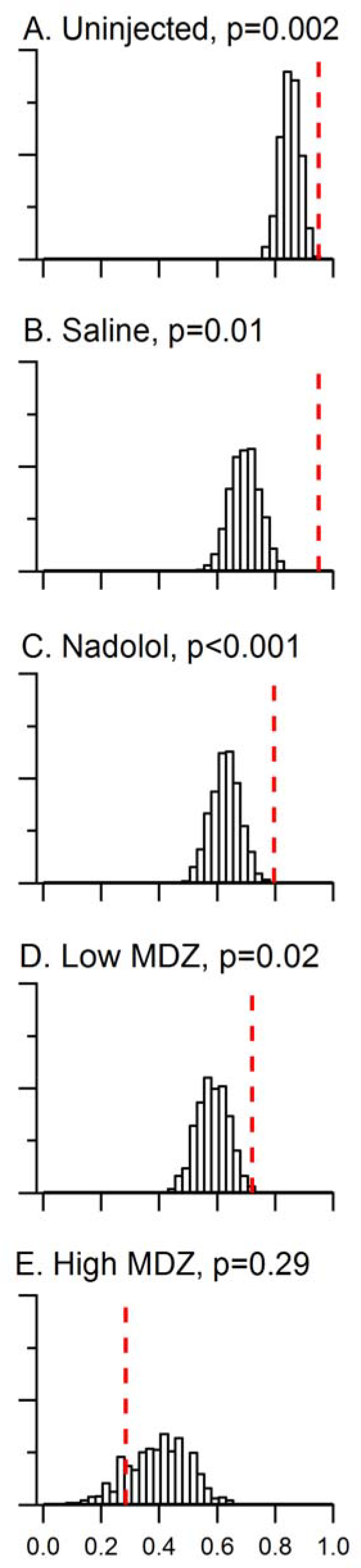
Day to day reinforcement occurred with a probability that was greater than chance for all groups in the trapped condition, except rats injected with the high dose of MDZ. A model that took into account the effects of learning and individual differences was constructed. The chance distribution of %SO (proportion of sequential openings) values from 10,000 matrices is shown in histogram form for each group tested with a trapped rat. The observed %SO is marked by the red dotted line in each panel and the two-tailed probability of the observed %SO occuring by chance listed. For uninjected rats or rats injected with saline, nadolol or the low dose of MDZ, the observed %SO was significantly greater than would be expected by chance. However in the case of rats injected with the high dose of MDZ, the observed %SO was less than 85% of the chance %SO values.

Several points are evident in comparing the observed and null distributions across groups. First, the proportion of reinforced openings predicted by chance was highest (median=0.85) and least variable (10 to 90 percentile range = 0.09) for uninjected rats. Second, for rats injected with the high dose of MDZ, the proportion of reinforced openings predicted by the null model was relatively low (median=0.38) and the distribution was broad (10 to 90 percentile range=0.30). Third and most importantly, the proportion of reinforced openings observed was always greater than the proportion of reinforced openings predicted by the null model for all groups except those treated with the high dose of MDZ. To quantify this comparison we calculated a conditional probability that the proportion of observed sequential openings occurred by chance (p(%SO|null), essentially a p-value). Lower values of p(%SO|null) reflect a greater likelihood that sequential day-openings did ***not*** occur by chance and are therefore a positive measure of the strength of day-to-day reinforcement. The p(%SO|null) was significant for rats that were either not injected or injected with saline, nadolol, or low MDZ; values ranged from <0.001 to 0.02, (Fig. 3A–Fig. 3D; see figure for p(%SO|null) values). Only in the case of rats injected with high MDZ was the observed pSO less than the median chance occurrence of reinforced opening; this group yielded a p(%SO|null) of 0.29 which was not significantly different from chance (Fig. 3E).

## Opening streaks

As would be expected as a result of reinforcement, past door-openings had a positive effect on the chance of a future door-opening. However because reinforcement differed across groups, so did the total number of sequential door-openings. We therefore analyzed streak length, meaning the number of sequential days that an individual rat opened the door. Uninjected rats were at one extreme with the highest number of sequential openings and rats treated with the high dose of MDZ were at the other extreme with the lowest number of sequential openings.

Uninjected rats opened on the day immediately following 78 of 81 openings that occurred on days 1-11 (96%). Furthermore, whenever an uninjected rat opened for two days in a row, he always opened on the next (third) day as well (69/69 opportunities). Because of this tendency to repeatedly open the restrainer door, uninjected rats opened for long streaks, including 2 animals that opened on all 12 days of testing (Fig. 4A). At the other extreme, rats treated with the high dose of MDZ opened the restrainer door on two sequential days on only 29% (4/14) of the opportunities and *none* opened for three days in a row (0/3 opportunities). Rats in the other groups opened for streaks of intermediate lengths (Fig. 4A).

**Figure 4.**
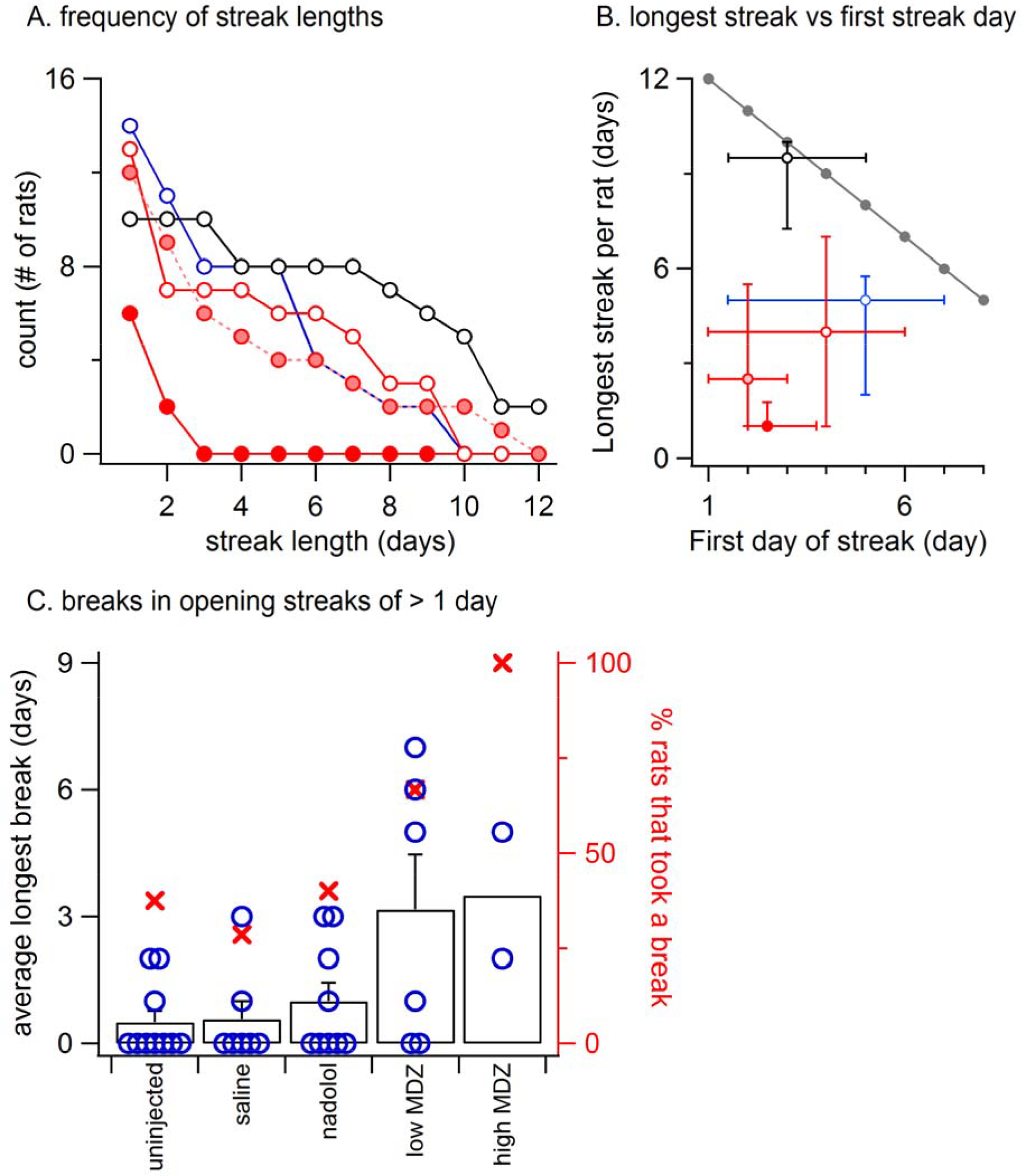
A: Using the same symbols as in Figure 1 (open black, uninjcted; open red, saline; pink, low MDZ; solid red, high MDZ), the frequency of opening streaks (consecutive day openings) of different lengths is illustrated for streaks of lengths from 2 to 12 days. At the left is the number of rats that opened at least once. B: Again, using the same symbols as in Figure 1, the median length of the longest streak (± 25 and 75 percentiles) is graphed as a function of the median testing day (± 25 and 75 percentiles) on which the streak began. The gray line at the top shows the optimal possible performance (e.g. rats that began opening on day 1 could achieve a streak of 12 days). C: The failure of a rat to open for one or more days is termed a “break.” An analysis of breaks for rats that opened on at least two consecutive days on days 9-12 (uninjected, n=10; saline, n=7; nadolol, n=8; low MDZ, n=6; high MDZ, n=2) shows that rats treated with MDZ were more likely to take at least one break (filled red circles, right axis). Rats treated with MDZ also took longer breaks on average than did rats from the other groups (black columns, left axis). The individual points for all rats considered in this analysis are illustrated by the hollow blue circles.

The maximal possible length of an opening streak is greatest when rats open on the first day and declines thereafter (Fig. 4B, gray line). We analyzed the longest streak for each rat and compared the average streak length (1-12 days) to the average first day of the streak (day 1-12) for each group of rats. For uninjected rats, the median first opening occurred on day 3. The median length of the opening streak by uninjected rats was nearly the maximum value of 9 days. In contrast, the opening streaks of rats from all other groups fell far short of the maximum possible.

It is notable that rats treated with MDZ started streaks earlier (day 1-5) than any other group but still had the shortest streak lengths (1-3 days). Thus the median streak length deviated from the theoretical maximal streak length by only 0.5 in the case of uninjected rats but by 9.5-10.5 days in MDZ-treated rats. The maximal streak length of nadolol- and saline-treated rats was less than the maximum possible by 3-5 days.

For rats that opened on at least two consecutive days on days 9-12 (uninjected, n=10; saline, n=7; nadolol, n=8; low MDZ, n=6; high MDZ, n=2), those treated with MDZ (either high or low dose) were more likely to take at least one break (red x-s, right axis of Fig. 4B) and also took longer breaks on average than rats from the other groups (black columns, left axis of Fig. 4C). This latter difference was significant between the 6 rats in the low MDZ group and the 10 rats in the uninjected group that met the criteria for this analysis (one-way ANOVA; F(4, 28)=3.81, p=0.01).

## Blocking distress in free rats tested with a chocolate-containing restrainer

To determine whether the reduction in door-opening observed in MDZ-treated rats could be due to an effect of MDZ other than its anxiolytic influence, such as sedation, rats were injected with a high (n=8) or low (n=8) dose of MDZ or saline (n=8) prior to testing with a restrainer containing chocolate, a non-social reward. As expected, the opening latency decreased across days (Fig. 5A). The decay in opening latency across days differed between treatment groups (χ^2^(2)=13.2; p=0.001). Rats treated with either dose of MDZ, but not those treated with saline, showed significantly decreasing opening latencies across the days of testing (saline: N(0,1)=0.5, p=0.62; low: N(0,1)=−4.30, p<0.001; high: N(0,1)=−3.67, p<0.001). On the final 3 days of testing, when latencies had plateaued, the learned latency was significantly different between groups (Fig. 5C; one-way ANOVA; F(2, 21)=3.955, p=0.04). Tukey post hoc tests revealed that the average opening latency in saline-treated rats was greater than in rats injected with the low dose of MDZ (p=0.04). As with saline-injected rats tested with a trapped rat, MDZ-treated rats tested with chocolate showed a shift from longer to shorter opening latencies across the days of testing (Fig. 2B). In contrast, saline-injected rats tested with chocolate did not show a shift in latencies across the days of testing (Fig. 2B).

**Figure 5.**
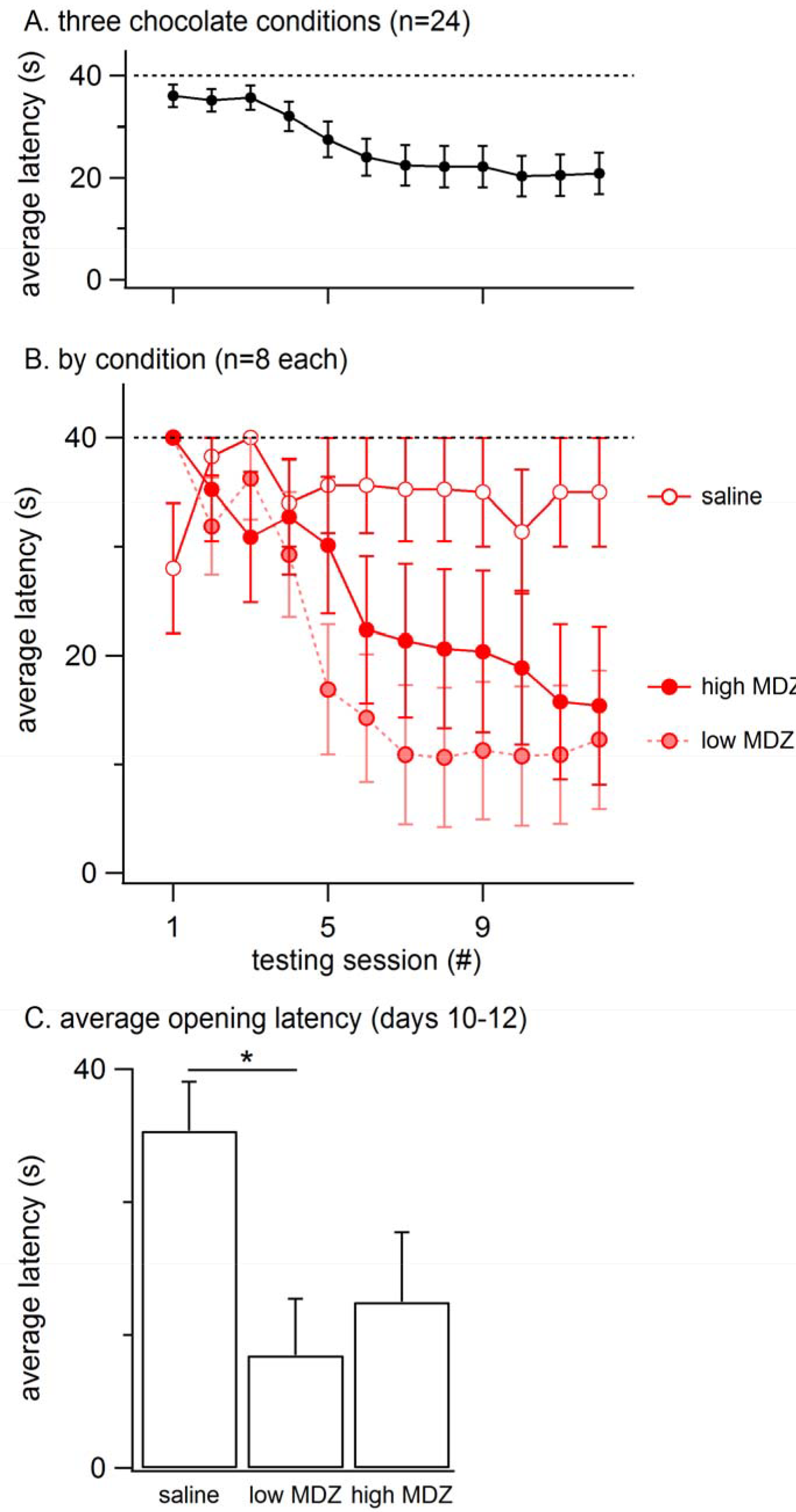
A: The mean (± SEM) latency to door-opening for all rats tested with a chocolate-containing restrainer (n=24) decreased across the 12 days of testing, suggestive of learning. B: The decay in door-opening latency across testing sessions differed between the groups of rats tested (n=8 per group). C: The average opening latency during the final three days of testing, at a time when latencies had stabilized, was significantly less for rats treated with a low dose of MDZ than for rats that received saline (*, p=0.04).

## Comparison of reinforcement in chocolate and trapped conditions

As introduced above, the strength of reinforcement was quantified as p(%SO|null), the conditional probability that the proportion of observed sequential openings occurred by chance. We performed this analysis for rats in both trapped and chocolate experiments. Yet the p(%SO|null) values from trapped and chocolate conditions could not be directly compared because of statistical power differences created by the different number of rats studied (trapped conditions N=16; chocolate conditions N=8). To enable a valid comparison, we reduced the statistical power of the trapped groups to the power level of chocolate groups. This was accomplished by a power-matched bootstrapping of the three trapped conditions that shared pharmacological manipulations with chocolate conditions (high MDZ, low MDZ, saline). For each trapped condition, we created 100 bootstrapped samples, each containing 8 rats randomly chosen from the 16 rats. Almost a quarter of the samples from the high MDZ-trapped condition (n=23) were removed due to the absence of any openings. Bootstrapped samples were then analyzed with the null model, thereby generating bootstrapped p(%SO|null) values (low MDZ N=100; Saline N=100; high MDZ N=77) that represented the strength of reinforcement in the trapped conditions *if they had been tested with the same statistical power as were the chocolate conditions*.

Since each bootstrapped p(%SO|null) from a trapped condition has the same power as each p(%SO|null) from a chocolate condition, we can compare strength of reinforcement between chocolate conditions and trapped conditions by comparing the p(%SO|null) values of a chocolate condition to the distribution of p(%SO|null) values of bootstrapped samples of the corresponding trapped condition (Fig. 6). As expected bootstrapped p(%SO|null) values (dashed line, saline: 0.08; low MDZ: 0.08; high MDZ: 0.85) were greater (farther from significance) than the original p(%SO|null) values (cross, 0.01, 0.01, 0.40), reflecting reduced statistical power.

**Figure 6.**
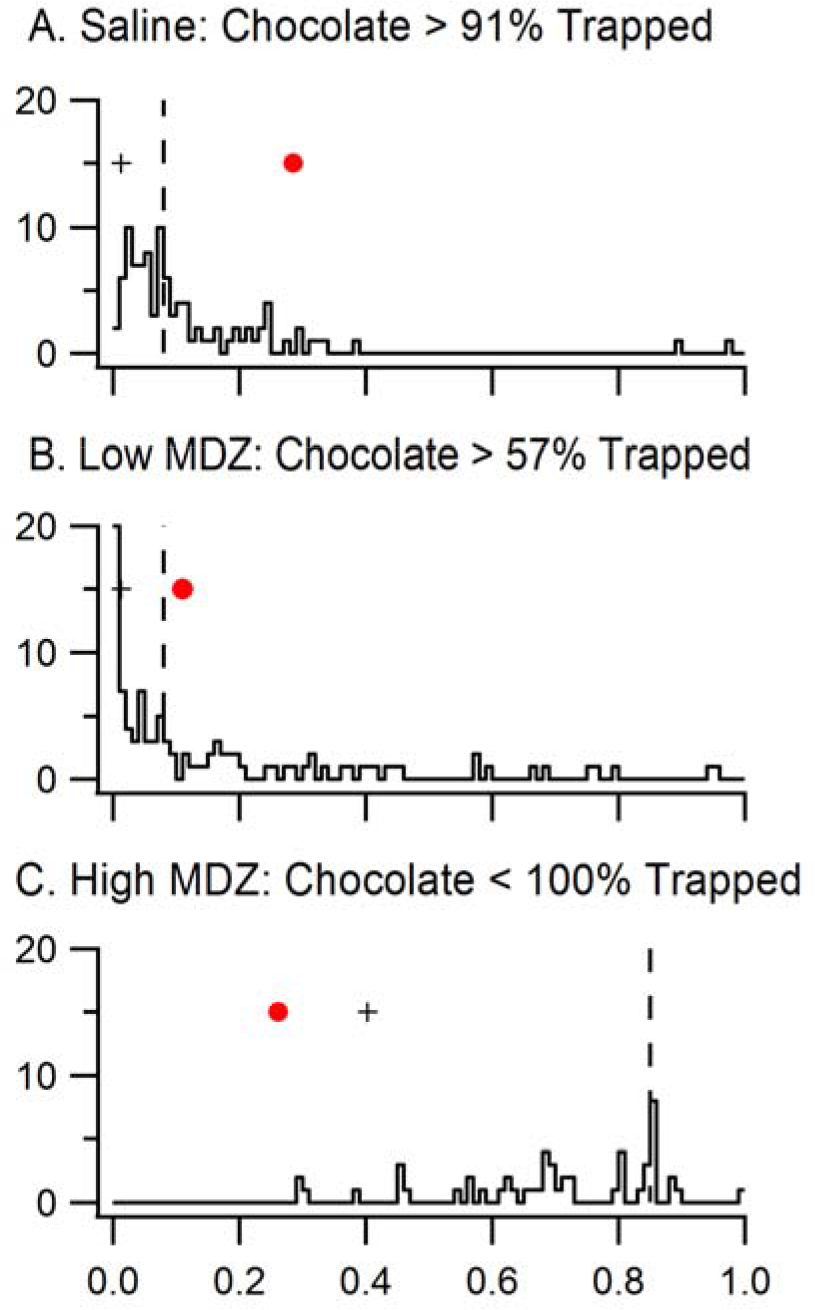
Data from rats tested with a trapped rat and injected with saline, low MDZ, or high MDZ was resampled to match the statistical power of chocolate conditions. The distributions of p(%SO|Null)s from 100 bootstrapped samples from each trapped rat condition are shown in the histograms. The medians of these bootstrapped p(%SO|Null)s (dashed line) are greater than the orginial p(%SO|Null) values (crosses) from the complete data set of trapped rat conditions, showing reduction in statistical power. The p(%SO|Null) of rats tested with chocolate (red dots) were compared to that of rats tested with a trapped rat.

The p(%SO|null) of the high MDZ chocolate group (red marker, 0.26) was significantly lower than the bootstrapped p(%SO|null) values of high MDZ-trapped (77 out of 77 bootstrapped values are higher than 0.26, p<0.01), reflecting that high MDZ-treated rats were more reinforced when chocolate, rather than a trapped rat, was in the restrainer. The p(%SO|null) of the low MDZ-treated rats tested with chocolate (red marker, 0.10) was higher than the bootstrapped P(%SO)s of low MDZ rats tested with a trapped rat (43/100 of bootstrapped values were lower than 0.10, p=0.43). This difference was not significant. This reflects a roughly equal strength of reinforcement between chocolate and a trapped rat for rats treated with low MDZ. Finally, the p(%SO|null) of saline-treated rats tested with chocolate (0.29) was significantly higher than the bootstrapped p(%SO|null) values from saline-treated rats tested with a trapped rat (9/100 were higher than 0.29, p=0.09), reflecting greater reinforcement by a trapped rat than by chocolate for rats treated with saline.

In sum, high MDZ treatment renders chocolate more reinforcing than a trapped rat whereas saline treatment renders the trapped rat more reinforcing than chocolate. For rats treated with low MDZ, the reinforcement engendered by a trapped rat and by chocolate were not different.

## Corticosterone responses to the helping behavior test

To determine the HPA reaction to vicarious distress, corticosterone (CORT) was measured in rats exposed to a trapped cagemate and compared to the CORT responses of the trapped rats themselves. CORT is an index of hypohalamo-pituitary-adrenal (HPA) axis involvement. In this experiment, CORT levels were measured following a 40- minute exposure to a trapped rat. The restrainer door was secured shut, ensuring that all free rats were exposed to the trapped rat for the full 40 minute duration. CORT responses were calculated by subtracting a pre-session baseline from a measurement taken immediately after testing (see Methods). After the experimental exposure to a trapped rat used to collect CORT, rats were tested in the standard paradigm described above for 12 days.

To test the relationship between CORT and helping behavior, a regression was performed between the average opening latency across the 12 days of standard testing and individual CORT responses. The CORT response of free rats was significantly correlated to the average opening latency across the 12 sessions (r^2^=0.52; F(1, 15)=16.54, p<0.001; Fig 7A). In contrast, no significant correlation existed between the CORT response of the trapped rats and the average opening latency (r^2^=−0.04; F(1,18)=0.69, p=0.42; Fig. 7B). Thus, a stronger HPA activation response is associated with little door-opening (high average latency) and individuals with weaker HPA reactivity opened the restrainer door at the lowest latencies.

**Figure 7.**
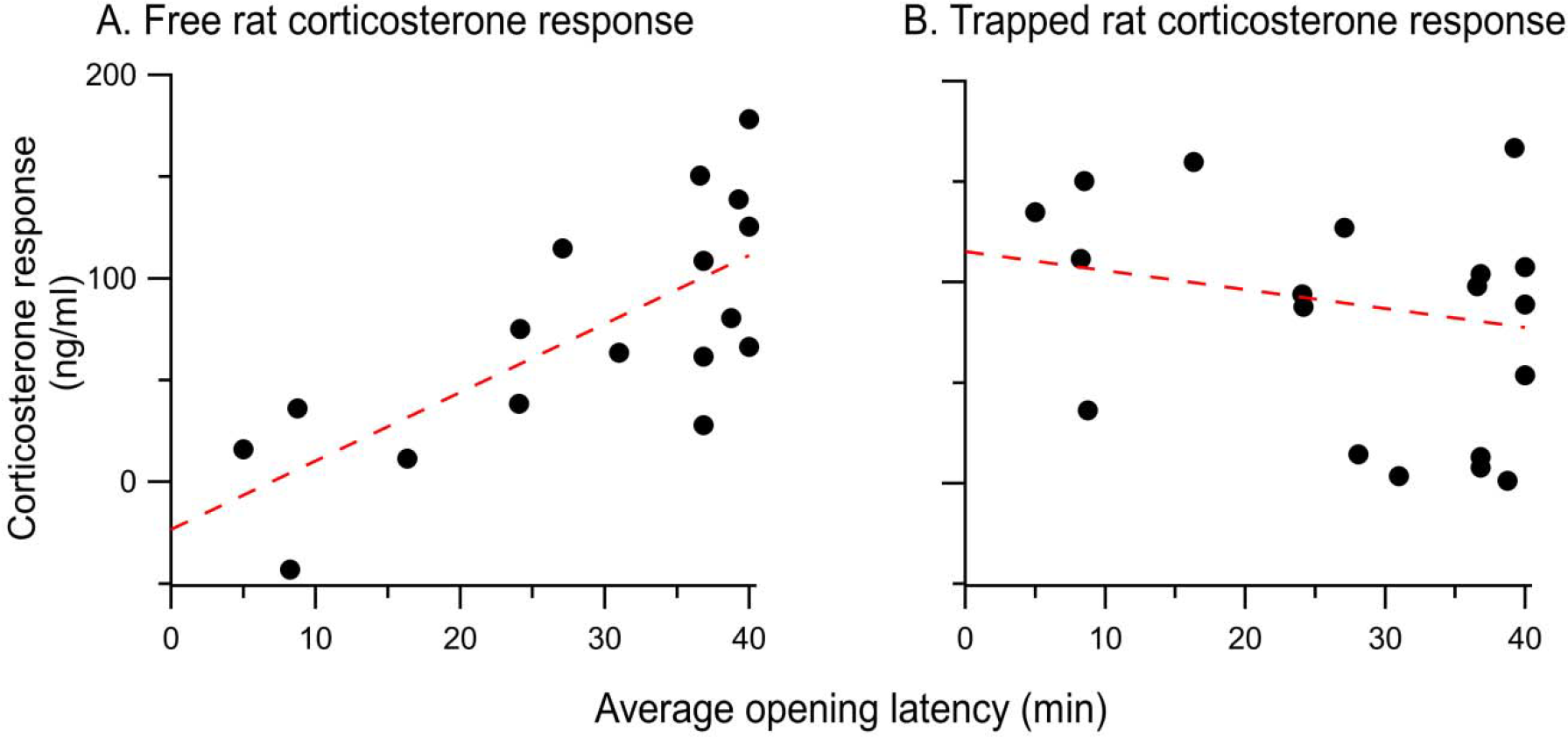
**A**: The corticosterone response of each free rat evoked by vicarious distress is positively correlated with the average opening latency over the 12 days of testing. behavior. **B**: There was no correlation between the the trapped rat’s corticosterone response and the ensuing opening behavior of the free rat.

## Discussion

This study demonstrates that the release of a trapped conspecific requires affective processing that is blocked by the benzodiazepine anxiolytic MDZ. Although rats treated with MDZ did not open a restrainer to release their trapped cagemate, they did open a restrainer to access chocolate. Thus, the reduction in pro-social behavior produced in MDZ-treated rats was not due to a sedative, cognitive, motor or non-specific and unidentified effect. Instead, MDZ interfered specifically with social affective processing that appears necessary to motivate a free rat to help a trapped rat. In humans, affective communication from one individual to another fuels an empathic understanding and pro-social actions. We hypothesize that a similar motivation underlies the rat’s action in the simple helping situation presented to them in the current experiments. Specifically these data support the idea that affective resonance between helper and victim rats is responsible for motivating pro-social actions.

While the idea that rats engage in emotionally driven social behaviors including helping behavior was originally received with hesitation, a recent explosion of studies confirms and extends this finding (Burkett et al., 2016; Hernandez-Lallement et al., 2015; 2016; Márquez et al., 2015; Muroy et. al., 2016; Sato et al., 2015). Of particular relevance here, two groups have established that rats favor pro-social (shared) food distribution over a selfish option. In these paradigms, actor rats receive food regardless of whether they provide another rat with food. Rats preferentially choose to provide food to another rat over receiving food alone (Hernandez-Lallement et al., 2015; Márquez et al 2015). The finding that rats choose to provide food to another, despite no added benefit conferred for doing so, is strong evidence that they are sensitive to the well-being of others. Interestingly, lesions of the amygdala block food-sharing (Hernandez-Lallement et al., 2016). Given the amygdala’s key role in affect and motivation, this result suggests that food-sharing in rats is affectively motivated, an interpretation that is consistent with the present findings. Thus converging evidence suggests that pro-social behavior in rats can be motivated by affect.

## Social interaction is not the motivation for helping

The failure of MDZ-treated rats to release a trapped cagemate is further evidence that rats are motivated by negative affect rather than by a desire for social interaction as has been recently argued (Silberberg et al., 2014). Animals motivated primarily by a desire for social interaction would have opened a restrainer containing a trapped rat just as they opened a restrainer to access chocolate. However, this did not happen. Therefore it appears that rats open only for a rat in distress and only when they are capable of mounting an affective response. This idea is in line with previous results. Rats repeatedly release cagemates even when subsequent social contact is prevented following their release (Ben-Ami Bartal et al., 2011). Moreover, in a recent experiment, the effects of distress were disambiguated from the effects of the size of the area in which a rat was trapped (Sato et al., 2015). Rats were trapped in a pool-arena filled with water. Rats in an adjacent arena of the same size opened the door to allow the soaked rats to access their dry arena. However, when rats were placed in the pool-arena *without water*, the rats did not open the door separating the two compartments. Thus rats only opened the door when a rat was distressed, strong evidence that the helper rats were not motivated by a desire for social contact. In sum, the opportunity to socially interact with a rat is neither necessary nor sufficient to motivate helping a trapped rodent in distress.

## A novel method for evaluating reinforcement

The method introduced here to quantify day-to-day reinforcement can be adapted for use in many experimental conditions. The prerequisites are a binary choice and sequential testing. The advantage to this method is that it is able to test whether the outcome of a previous decision positively or negatively reinforces the decision while removing confounding effects created by non-associative learning that takes place across sessions. It therefore tests the strength of reinforcement against a conservative null hypothesis and quantitatively represents the strength of reinforcement.

## The relative motivational value of helping and chocolate

We found that saline treatment reduced the motivational value of accessing chocolate below the motivational value of opening the restrainer door for a trapped cagemate. At first glance, these results suggest that helping may be more highly valued than chocolate, particularly in the anxiogenic conditions created by receiving an injection. An alternative possibility stems from the *social buffering* afforded by the presence of two rats in the trapped condition and only one rat in the chocolate condition. *Social buffering* refers to the anxiolytic or emboldening effects afforded by the presence of a conspecific (Kikusui et al., 2006). Thus, it is possible that social buffering effectively allowed saline-treated animals to enter the arena center and open the door to a restrainer containing a cagemate. Yet this interpretation is hard to reconcile with the findings that MDZ-treated rats did not open the restrainer despite the presence of the trapped cagemate. In other words, even though MDZ-treated and saline-treated rats tested with a trapped cagemate enjoyed the same social buffering, only the saline-treated rats ventured into the arena center and helped the cagemate by opening the restrainer door. Thus social buffering cannot be at the root of the behavioral differences observed between the two groups. Moreover the pharmacological reduction in anxiety produced by MDZ would exert an influence similar to social buffering and also cannot form the basis for the group differences.

The most parsimonious explanation is that MDZ treatment antagonizes the motivation to release the trapped rat through blunting the affective processing of social cues emanating from the trapped rat. Under this scenario, a reduction in anxiety can help facilitate entry into the arena center but is insufficient without a strong source of motivation. In the trapped rat condition, the source of motivation is the affect evoked in the free rat by the trapped rat. The high dose of MDZ effectively neutralized this affective motivation and thereby blocked helping.

Rats injected with saline did not access chocolate although MDZ-treated animals did. This result may stem from an increase in food palatability that has been reported after MDZ treatment (Gray and Cooper, 1995). Another possibility (that is not mutually exclusive) is that the anxiogenic effects of the injection procedure antagonize opening. Saline treatment is pharmacologically inert but nonetheless induces stress even in habituated rats. In this light, the ability of saline-treated rats to overcome their anxiety and help a trapped rat demonstrates their degree of motivation, and puts into stark relief the complete lack of motivation observed in the non-stressed, MDZ-treated rats to release their conspecifics.

## MDZ blocks helping through central actions

The MDZ experiments reveal that anxiolysis blocks helping behavior, but do not unequivocally establish the physiological mechanism for this effect. In order to determine the role of sympathetic arousal in motivating helping, we tested the peripherally acting beta-adrenergic blocker, nadolol, which blocks sympathetic activation but does not cross the blood-brain barrier and therefore leaves central affective circuits and HPA activity unaltered. Nadolol treatment had no effect on helping behavior, resembling a saline injection in all respects. This result shows that MDZ does not block helping through a sympatholytic effect. The finding that rats with the smallest corticosterone responses to viewing a trapped rat were the best helpers suggests that the effects of MDZ are not due to the indirect effect. Rather, it is likely that MDZ acts to block helping through central actions on affective circuits.

## The effect of stress on helping behavior follows an inverted U-shaped curve

Our results suggest that the effect of stress on pro-social behavior follows an inverted U-shaped curve, making moderate levels of stress most conducive to helping. The finding that MDZ treatment blocked helping supports the idea that low levels of negative arousal reduce the motivation to act for the benefit of another rat. On the other end of the spectrum, rats that had a high CORT response upon exposure to a trapped cagemate were less likely to help than were rats that showed smaller CORT responses. Together these findings suggest that both low and high levels of negative arousal are detrimental to successful helping. This relationship is similar to the effects of stress on learning, memory and performance tasks wherein a moderate stress response enhances performance, while too little or too much stress is detrimental (Joëls 2006, Salehi et al 2010, Sapolsky 2015, Schilling et al 2013). Muroy et al. (2016) recently reached a similar conclusion for the effect of stress on pro-social behavior. They found that moderate stress (restraint) increased water-sharing whereas extreme stress (restraint paired with predator odor) reduced sharing in rats.

Others have also found that pro-social behavior is negatively impacted by HPA activity in humans and other animals. In humans, personal distress opposes the expression of other-oriented empathy (Batson et al., 1987). Individuals with the short allele polymorphism of the serotonin transporter gene regulatory region (5-HTTLPR) have higher HPA reactivity (Gotlib et al., 2008) and lower pro-social tendencies (Stoltenberg et al., 2013). Physiological stress as measured by HPA reactivity therefore appears to antagonize helping, rendering individuals “afraid to help” in the words of Stoltenberg and colleagues. In chimpanzees, increased HPA reactivity is associated with a reduced propensity for pro-social behavior (Clay and de Waal, 2013).

Consistent with our finding that helping is negatively correlated with large HPA responses to the distress of another, administration of a glucorticoid synthesis inhibitor extends empathic responses to strangers in mice and humans (Martin et al., 2015). In the social prairie vole, observer animals increased their grooming of demonstrator conspecifics that had been shocked during a separation (Burkett et al., 2016). Since this increase in other-oriented grooming behavior did not occur during reunions with naïve (not shocked) animals, the behavior was interpreted as representative of consolation. Of great interest, the corticosterone responses measured from observer and demonstrator voles were strongly correlated when the demonstrator was shocked but not when he was naive. This result shows that shocked voles vicariously communicate their distress to observer conspecifics. Thus, emotional contagion between voles is expressed through HPA state-matching, an idea that is supported by work on emotional contagion for pain in mice and humans (Martin et al., 2015). Yet, it is not clear whether the prosocial behavior of consolation (allogrooming) also correlated with the demonstrator’s corticosterone response. Consequently, the result cannot be directly compared to our and other’s findings on the effect of HPA responses on pro-social behavior.

It may appear paradoxical that blocking anxiety through MDZ treatment prevents rats from helping whereas low HPA reactivity appears to allow or possibly promote helping. It is worth stating that MDZ-evoked anxiolysis is not synonymous with a low HPA response. CORT exerts a complex influence on social behavior through effects on central pathways and the effects of CORT strongly interact with trait anxiety in rodents (Beery and Kaufer, 2015). The affect of anxiety, a central emotional state, is only one of many influences on HPA activity, which is notably increased by positive as well as negative arousal. It remains unclear to what extent the HPA responses measured in the present study directly cause a decrease in helping or are simply an index measure of the causative agent.

## Conclusion

In conclusion, this series of experiments clearly demonstrates the fundamental role of affective arousal in motivating rats to help their cagemate escape a trapping restrainer. The helping behavior shown by rats in the present study is not a conditioned response motivated by either approach to a positive reward or avoidance of a negative cue. Rather, the data presented here support the idea that rats resonate with the negative arousal of the trapped cagemate and are moved to approach the cagemate because of this affective response. As in humans, rats find helping rewarding, as witnessed by the recurrence of door-opening on sequential days. The benefit of door-opening for a trapped rat can be quantified and compared to door-opening for chocolate, paving the path for studies considering the cost and benefit of pro-social behaviors. Finally, a moderate level of arousal was the best predictor of pro-social behavior. This suggests that pro-sociality in rats has an inverted U-shape relationship to stress.

The rats’ response to a trapped cagemate shares multiple elements in common with empathically motivated helping behavior in humans, and can serve as a model for studying the biological mechanisms of human pro-sociality.

## Author and contributors

Trapped rat experiments: design (IB, PM, JD), data acquisition (IB, HS, NM, TM, PM); Chocolate experiments: design (IB, PM), data acquisition (IB, JW); Corticosterone experiments: design (IB, PM), data acquisition (IB, PM); Data analysis & statistics: (IB, HS, NM, TM, PM); Reinforcement model: (HS, PM); Drafting and revising manuscript: (IB, HS, NM, TM, JW, JD, PM).

## Acknowledgements

The assistance of Miguel Barajas, Isabel Boni, Tony Logli, Maria Sol Bernardez-Sarria, Katie Ragsdale, David Rodgers, Yuri Sugano, and Jenny Wang, is gratefully acknowledged. We’d like to thank Dr. David White and Fanny Delebeque for their help with acquisition of corticosterone measurements. The authors are indebted to Peter McCullagh for patient and expert advice on crafting and coding the general linear models used.

